# OST component RPN1 is a novel regulator of IRE1 RNase activity that interacts with multiple distinct IRE1 and PERK complexes

**DOI:** 10.1101/2024.12.17.628858

**Authors:** Natacha Larburu, Freya Storer, Yassin Ben-Khoud, Chao-Sheng Chen, Christopher J Adams, Tony D Southall, Maruf MU Ali

**Author notes:** Present address: MSA sud Aquitaine place, Marguerite Laborde, 64017 PAU, France. Present address: Department of Medicine, Imperial College London, London SW7 2AZ, UK. Present address: UCB Pharma Ltd, 216 Bath Rd, Slough, SL1 3WE. These authors contributed equally to the work (Natacha Larburu, Freya Storer).

## Abstract

The unfolded protein response (UPR) is an essential cell signalling system that regulates ER protein homeostasis. IRE1 and PERK are receptor proteins that propagate the UPR signal from the ER to the cytosol. Both receptors are suggested to interact with various proteins from different biological pathways, although the scale and scope of such interactions are unclear. Previous reconstitution experiments have utilized purified isolated domains of IRE1 and PERK to understand mechanism. Here, we affinity purify full length IRE1 and PERK from mammalian cells and characterise the complexes they form by biochemical techniques and assess RNase function in vivo. We identify RPN1 as a novel interacting protein present in complexes with IRE1 and PERK. In the Drosophila eye, RNAi knockdown of RPN1 results in loss of IRE1 RNase activity. This work provides a basis for understanding of protein interaction networks for IRE1 and PERK and identifies OST subunit RPN1 as a novel regulator of IRE1 RNase activity.

## Introduction

The endoplasmic reticulum (ER) is the first compartment of the secretory pathway and plays an essential role in folding and maturation of membrane and secretory proteins within the eukaryotic cell. In certain situations, the influx of nascent proteins into the ER can overwhelm the ER folding capacity resulting in an accumulation of non-native and misfolded proteins that are deleterious for cell fitness. The unfolded protein response (UPR) is a cell signalling system that detects misfolded proteins and elicits a cellular response that aims to restore protein homeostasis within the ER. This response includes transcriptional upregulation of UPR target genes, general cell translational attenuation, induction of ER associated degradation pathways and an increase in the size of the ER^1,2^.

IRE1 and PERK are two UPR activator proteins that either directly sense ER stress or indirectly via the Hsp70 ER chaperone BiP^3–5^. Both IRE1 and PERK share significant structural similarity within their luminal domains and have cytosolic kinase domains that further propagate the signal^6–8^. IRE1 stimulation leads to trans autophosphorylation and oligomerisation which in turn activates IRE1 C-terminal endonuclease to splice XBP1 mRNA^9–12^. This results in the formation of sXBP1 mRNA, which codes for a potent transcriptional factor that upregulates UPR target genes. For PERK, autophosphorylation in trans results in phosphorylation of eIF2α and inhibition of ribosomes to attenuate global cell translation.

IRE1 interaction with BiP has been well studied and is thought to be important in acting as an ER stress sensor^13–16^ and inhibiting the UPR signal^5,17^. Interestingly, emerging data indicates that IRE1 can interact with a variety of proteins from several different biological pathways and systems^18,19^. This includes a high affinity association with ribosomes^20^ and components of the protein translocation machinery including SEC61 translocon^21^ and SEC63^22^. These associations are consistent with a role in which XBP1 mRNA-ribosome-nascent chain complex is recruited to the ER membrane where an XBP1 dependent translational pausing mechanism enables productive interactions with IRE1 facilitating the splicing of uXBP1 to form sXBP1^23^. In addition, IRE1 can bind to other factors including chaperones Hsp47^24^ and PDIA6^25^, as well as influencing calcium signalling^26^ and DNA damage response within cells^27^. Much less is known about what interactions PERK may form besides its association with BiP^28^ and eIF2α^11^. Thus, understanding the interactions mediated by IRE1 and PERK, whether they form distinct complexes with binding partners, and what constitutes such complexes, is important to gain insights into possible connections between UPR and other biological systems.

Many previous studies of IRE1 and PERK have utilised isolated luminal or cytosolic domains in a reconstituted system to generate understanding of mechanism. This strategy has yielded great success but is limited to certain regions of the protein and does not cover the complete receptor. Using full length proteins that encompass the luminal domain, transmembrane and the cytosolic region, would provide much greater understanding to how the different domains communicate and interact with partner proteins, but has been a difficult technical hurdle to overcome.

In this study, we use native gel, mass spectrometry, and western blot analysis to characterise the interactions and complexes mediated by full length IRE1 and PERK proteins that are FLAG affinity purified from Expi293F cells. We find that the IRE1 and PERK form multiple distinct ER complexes, with several interacting partner proteins. Furthermore, we identify RPN1, a component of oligosaccharyl transferase complex (OST), as being an integral part of those complexes. In Drosophila, we introduce *RPN1* RNAi knockdown into transgenic flies that show reduced levels of IRE1 RNase activity. This current study provides a foundation for further analysis of protein interaction networks for IRE1 and PERK and suggests RPN1 as a novel effector of IRE1 splicing.

## Results

Protein expression constructs of IRE1 and PERK consist of their native N-terminal signal sequence followed by a triple FLAG tag and then the mature full-length protein sequence (Figure 1A). The generated insert was subsequently cloned into a pcDNA3.1 vector backbone. The constructs were expressed in Expi293F suspension cells, which can support a large volume of cell growth, thus generating sufficient recombinant protein for in vitro experiments (Figure 1B). The Expi293F cells were harvested and clarified, and the microsome/membrane fraction was resolubilised with detergent GDN101, a synthetic version of digitonin. IRE1 and PERK proteins were purified from cell lysate by FLAG affinity chromatography and were biochemically assessed. SDS-PAGE analysis indicates large over expressed bands at ∼130kDa and ∼150kDa that correspond to the predicted size of full length IRE1 and PERK proteins (Figure 1C,D). To confirm the identity of bands within the SDS PAGE gel, the bands were excised, trypsin digested, and the isolated peptides were subjected to Electrospray ionisation tandem mass spectrometry (ESI MS/MS) performed at the mass spectrometry facility (St Andrews University). The generated spectra were searched against sequence databases using the Mascot server^29^ (Supplementary Data 1). ESI MS/MS analysis confirmed the bands at 130kDA and 150kDA were IRE1 and PERK, thus demonstrating that we successfully purified recombinant full length receptor proteins.

**Figure 1.**
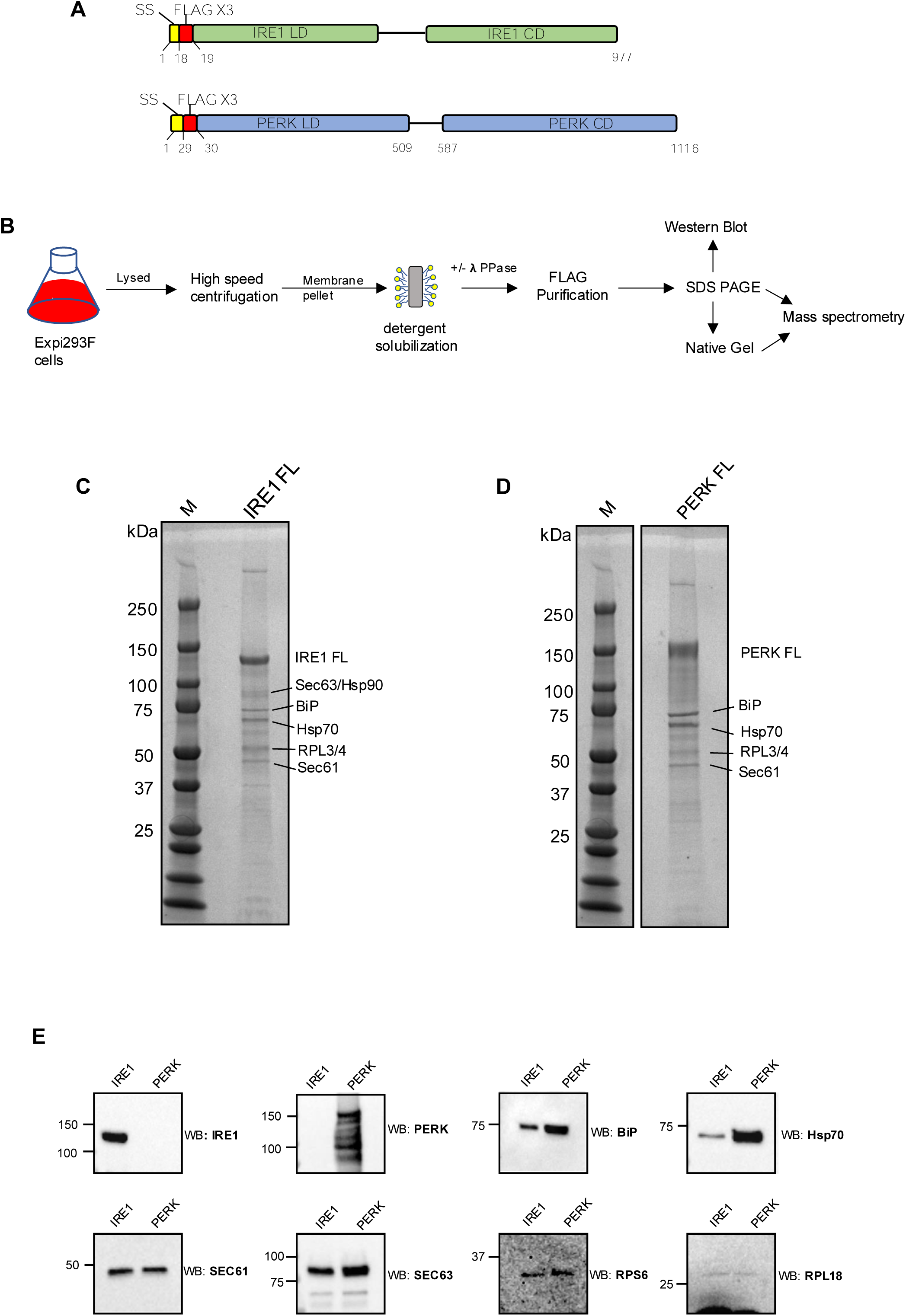
FLAG affinity purification of full length IRE1 and PERK. a) A drawing showing the full-length constructs that were expressed in Expi293F cells for IRE1 and PERK. The N-terminal native signal sequence for both proteins was followed by a triple FLAG tag and then the mature protein sequence encompassing the luminal domain (LD), linker and transmembrane region, followed by the cytosolic domain. B) an illustration showing the experimental workflow. C) SDS PAGE gel showing FLAG affinity purified IRE1 with over expressed band at 130kDa corresponding to full length protein. Other significant bands listed on the gel were identified by mass spectrometry analysis (Supplementary Data 1). The gel image is representative of 5 independent experiments. D) same as C but with over expressed band of ∼150kDa corresponding to full length PERK. E) Western blot analysis of selected proteins from both IRE1 and PERK affinity purified samples.

There were several other bands present along with both IRE1 and PERK suggesting that these receptors interact with other proteins from cell lysate. The other significant bands were identified as Hsp70 (73kDa) and BiP (70kDa), SEC63/Hsp90 at ∼90kDA, ribosomal proteins RL3/4 and OST at band around ∼50kDa, SEC61 at 37kDa. The appearance of a ladder of proteins below 50kDa is also suggestive of ribosome association (Figure 1C,D). The binding with ribosomes, SEC61 and SEC63 have been previously characterised for IRE1^20,21^, however, this data indicates PERK may also mediate similar interactions with protein translocation components.

As a secondary measure to assess protein interactions with IRE1 and PERK, we analysed the purified sample by western blot. We targeted proteins that were suggested by the mass spectrometry analysis and from previous studies that showed interactions with translocation components including, ribosomes^20^, SEC61^21^ and, SEC63^22^. To detect ribosomal proteins, we used RPS6 and RPL18 antibodies from both the small and large subunit of the ribosome. The resulting western blots showed a clear signal for SEC61, SEC63, RPL18, RPS6, Hsp70 from both FLAG purified IRE1 and PERK samples (Figure 1E), thus reinforcing the interactions between protein translocation components and IRE1 and PERK.

Although SDS PAGE analysis of FLAG purified IRE1 and PERK provide information about binding interactions between proteins, we wanted to assess the formation of distinct IRE1 and PERK complexes. For this purpose, we decided to use blue native gel electrophoresis. We collected the IRE1 and PERK sample after FLAG purification and subjected it to native PAGE and observed the signal produced from the gel by Coomassie staining. The native gel of IRE1 and PERK indicate about ∼6 distinct bands with both proteins displaying similar profiles (Figure 2 A,B). To identify what proteins constitute each distinct band from the native gel, we excised the bands and performed analysis by ESI MS/MS.

**Figure 2.**
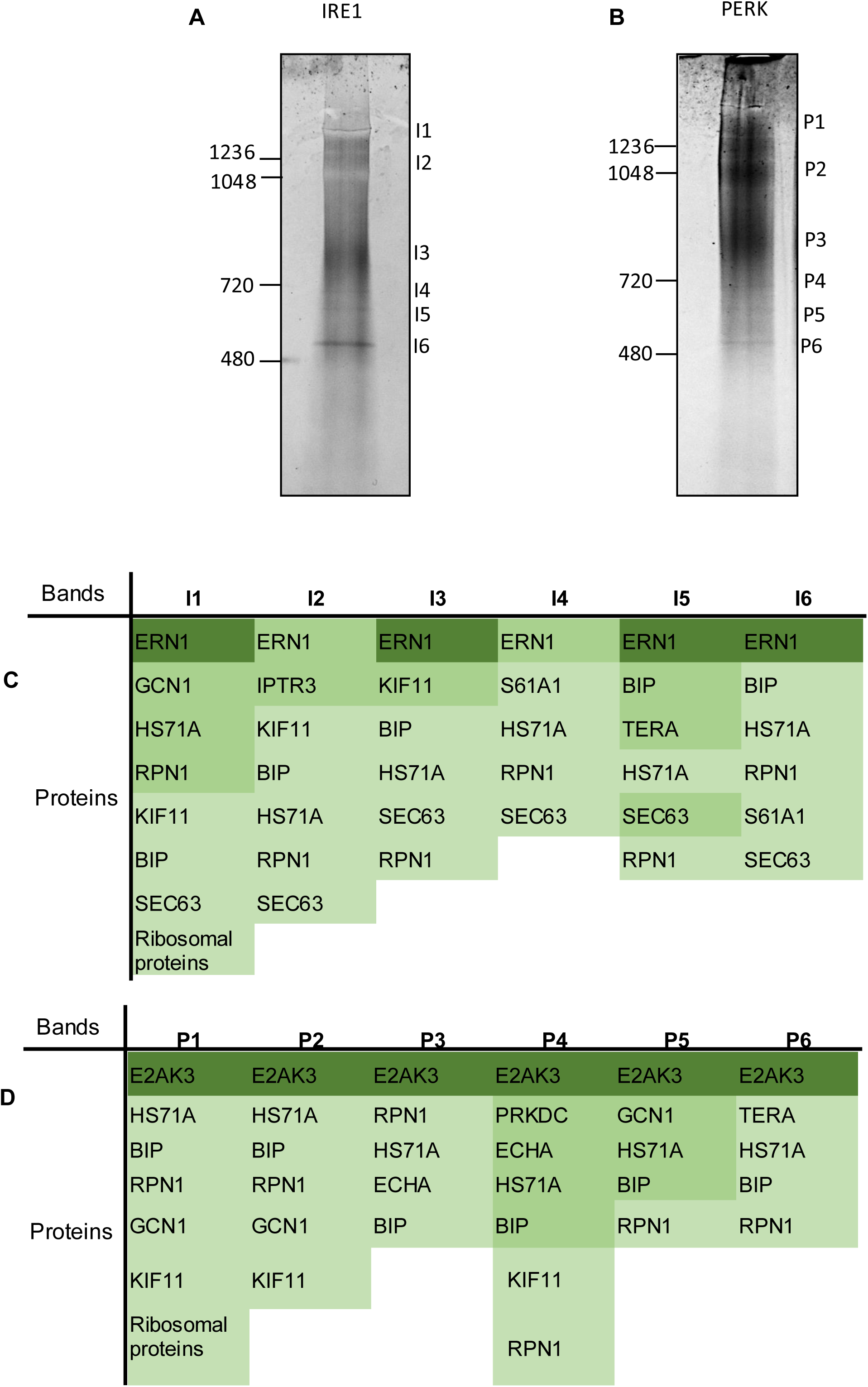
Native gel analysis of IRE1 and PERK. A) FLAG affinity purified full-length IRE1 sample was run on 3-12% Bis-Tris native gel and visualised with G-250 dye with protein standards listed to indicate size. There are 6 bands of varying resolution that suggest the formation of distinct IRE1 complexes. The gel image is representative of 3 independent experiments. B) same as A but for PERK. C) A list of selected proteins identified by LCMS/MS in order of their Mascot protein score for each of the bands I1-I6 identified in the native gel for IRE1. The proteins are coloured deep green for protein score of >2000, mid green >1000, light green >150. A protein score of >35 indicates identity or extensive homology. D) same as C but for PERK bands P1-P6.

Analysis of the mass spectrometry results were based on the Mascot protein score, which is the summed score for individual peptides that match a particular protein and the relative abundance of the protein within the sample as derived from the emPAI value (exponentially modified protein abundance index). The first band for both IRE1 (I1) and PERK (P1) is positioned above the 1236kDa marker. Our interpretation of the data indicates that this band forms a complex that contains IRE1, GCN1, Hsp70, RPN1, KIF11, BIP, SEC63 and several ribosomal proteins including RL6, RL7, RL10A, suggesting that this high molecular weight IRE1 complex interacts with ribosomes. A similar result is obtained with PERK, but without SEC63 (Figure 2 C,D; Supplementary Data 2 and 3).

The next band for both IRE1 (I2) and PERK (P2) is between the 1236 and 1048 kDa maker. For I2, the highest protein score besides IRE1 is for ITPR3 followed by KIF11, BiP, Hsp70, RPN1 and SEC63. However, ITPR3 has a low relative abundance compared to IRE1 and other interacting proteins. P2 consists of the same proteins as the equivalent IRE1 band without ITPR3 and SEC63, but with the addition of GCN1. A band appears between 1048 and 720 kDa for both IRE1(I3) and PERK (P3). These bands contain RPN1, Hsp70, BiP, for each receptor and additionally KIF11 and SEC3 for IRE1, whereas ECHA is observed for PERK specifically. The next three bands for IRE1 (I4-I6) and PERK (P4-P6) range from below 720kDa marker to above the 480kDa marker and are highly resolved forming narrow distinct bands. They contain for both IRE1 and PERK: Hsp70, RPN1, BiP, with SEC61 and TERA present in I5 and I6, and PRKDC, ECHA and KIF11 present in P4 with TERA present in P6. PRKDC has a high protein score but a low relative abundance when compared to PERK in P4 band (Figure 2; Supplementary Data 2 and 3).

Analysis of the different proteins identified in the various bands for IRE1 and PERK suggests a core complex of proteins that seem to associate with the receptors that consist of RPN1, Hsp70, BiP, with the addition of SEC63 for IRE1. Furthermore, this data newly identifies RPN1, a component of the OST complex, as an interacting partner present in IRE1 and PERK complexes.

Next, we wanted to measure the effect of ER stress on the formation of IRE1 and PERK complexes. We initially challenged Expi293F cells expressing IRE1 and PERK with tunicamycin and then compared the levels of phosphorylation for IRE1 and PERK to cells without tunicamycin induced ER stress. We observed that the levels of IRE1 and PERK phosphorylation were similarly high for both cells with and without the addition of tunicamycin (Supplementary Data 4). This indicates that expressing our protein in Expi293F suspension cells subjected the cells to a high degree of ER stress, so much so that chemical induction using tunicamycin had no further significant increase in phosphorylation levels.

In order to compare our ER stressed sample to a non-stressed sample we added lambda phosphatase to the cell lysate prior to purifying by FLAG affinity chromatography. Using this method would enable a comparison of our protein generated on ER stress to an inactive dephosphorylated IRE1 and PERK that could mimic a non-stressed sample.

The addition of lambda phosphatase almost completely dephosphorylated both IRE1 and PERK. This suggests that we have successfully generated both an ER stressed and active IRE1 and PERK, and an inactive dephosphorylated sample for comparison. Native gel analysis showed little difference between IRE1 and PERK with and without lamda phosphatase treatment (Figure 3 and Figure 4). We then attempted to measure the interacting proteins identified from the native gel and mass spectrometry analysis, by western blot. Initially, western blots were taken directly from native gel but displayed a smeared signal making interpretation very difficult. Thus, we compared IRE1 (Figure 3) and PERK (Figure 4) with and out treatment of lambda phosphatase by western blotting samples derived from the SDS-PAGE gel after FLAG affinity purification. We tested for IRE1 and PERK; the UPR sensor/inhibitor protein BiP; components of protein translocation, SEC61, SEC63, GCN1; ribosomal proteins, RSP6, RPL18; chaperone protein Hsp70, and other proteins including ITPR3 and RPN1 that were suggested by the mass spectrometry analysis. For each of the interaction partner proteins analysed there were no significant differences in binding when comparing the ER stressed IRE1 and PERK to the dephosphorylated sample which had been treated with lamda phosphatase. The only exception was for SEC63 where there was noticeably less SEC63 bound to IRE1 in the absence of lamda phosphatase (Figure 3C). Similarly, this was observed for PERK, but the result was more pronounced (Figure 4C). Thus, the interaction of partner proteins within complexes maybe independent of IRE1 and PERK phosphorylation activity.

**Figure 3.**
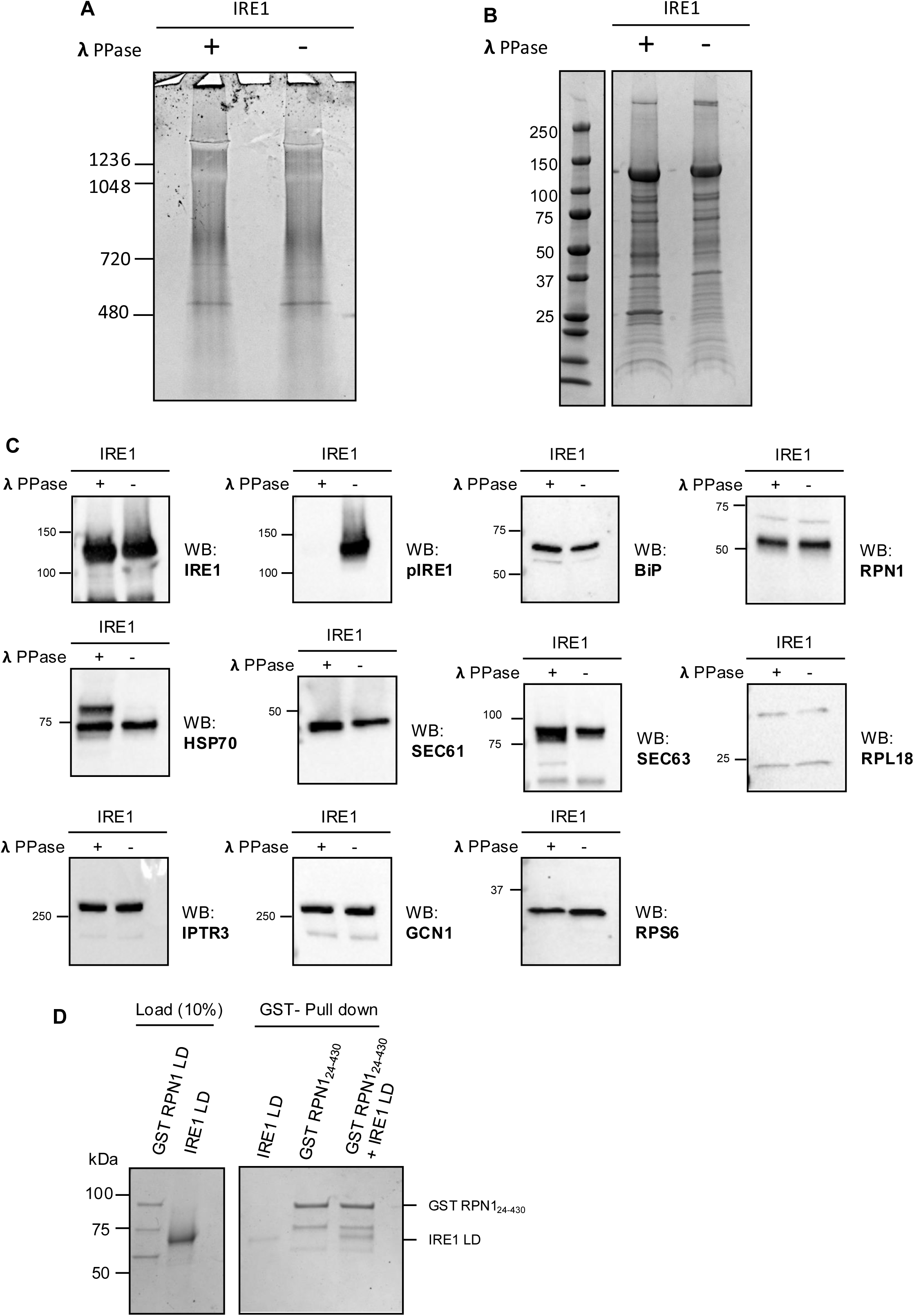
Analysis of affinity purified full length IRE1 subjected to **λ**PPase. A) Detergent solubilized IRE1 was incubated with **λ**PPase and FLAG resin for 1hr at RT with gentle agitation. The sample was subsequently FLAG affinity purified and run on 3-12% Bis-Tris native gel. This was compared to IRE1 sample without **λ**PPase treatment. B) Denaturing SDS PAGE gel of IRE1 with and without **λ**PPase treatment. The bands at 27 and ∼50 kDa are **λ**PPase and a degradation product. C) Western blot analysis of selected proteins performed on SDS PAGE IRE1 sample with and without **λ**PPase treatment. Images from A, B, C, are representative of 3 independent experiments. D) Pull-down assay using GST tagged RPN1 luminal domain (24-430aa) as bait protein, with IRE1 LD protein as the prey. Samples were run on an SDS PAGE gel stained with Coomassie and is representative of two independent repeats.

**Figure 4.**
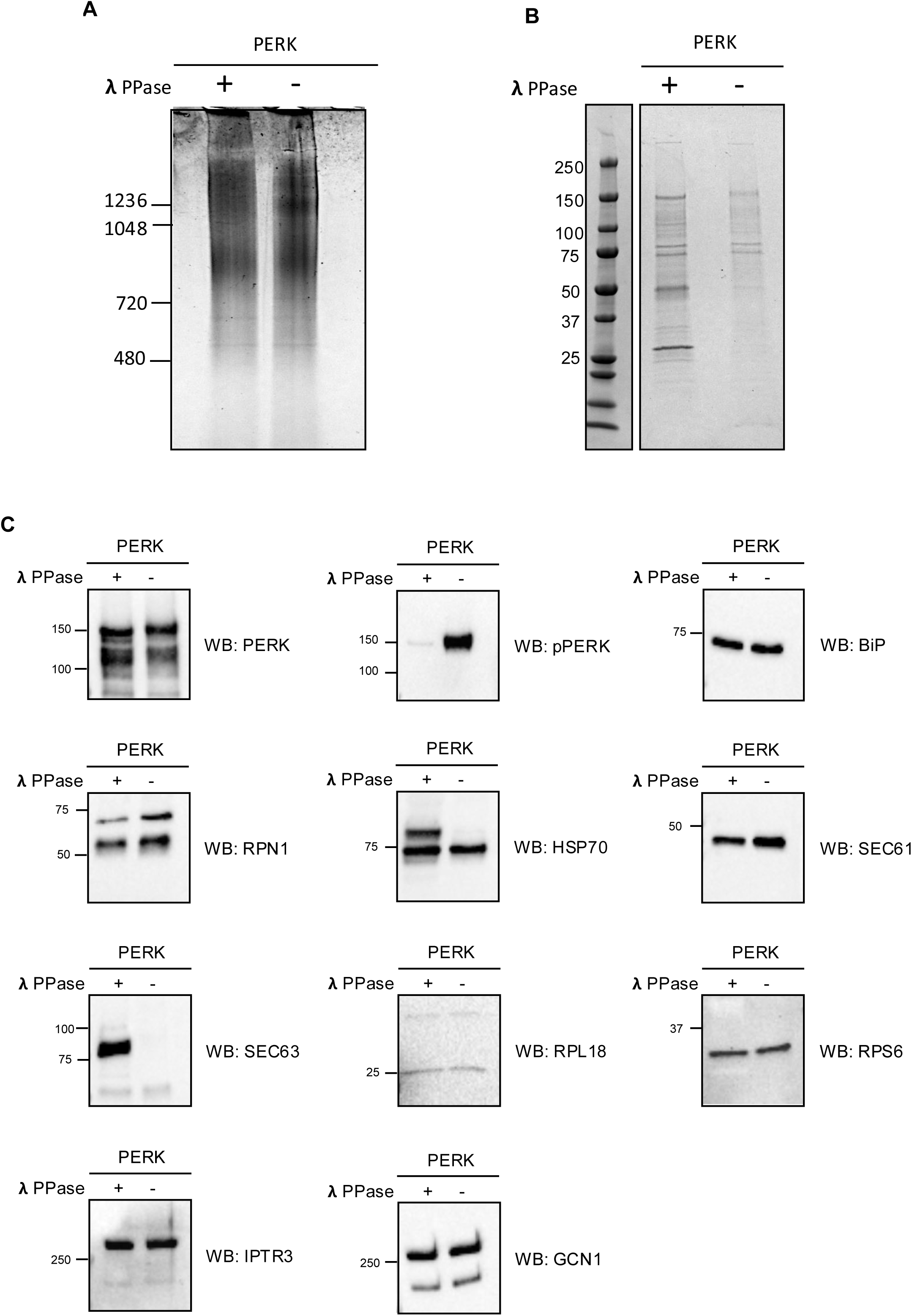
Analysis of affinity purified full length PERK subjected to **λ**PPase. A) Detergent solubilized PERK was incubated with **λ**PPase and FLAG resin for 1hr at RT with gentle agitation. The sample was subsequently FLAG affinity purified and run on 3-12% Bis-Tris native gel. This was compared to PERK sample without **λ**PPase treatment. B) Denaturing SDS PAGE gel of PERK with and without **λ**PPase treatment. The bands at 27 and ∼50 kDa are **λ**PPase and a degradation product. C) Western blot analysis of selected proteins performed on SDS PAGE PERK sample with and without **λ**PPase treatment. Images from A, B, C, are representative of 3 independent experiments.

Although we identify RPN1 in complex with IRE1 from the native gel, we did not know whether this interaction occurred directly with IRE1 or indirectly. We generated a construct of RPN1 that consisted of the full luminal domain, which encompasses the majority of the protein. The large size of the RPN1 luminal domain relative to the rest of protein structure suggested to us that this would be the more likely site of interaction with IRE1 via it’s LD. We conducted a pull-down assay to test for an interaction between the two proteins (Figure 3D). We observed an IRE1 LD band associating with the GST-RPN1 LD in a pull-down experiment. This suggests that IRE1 and RPN1 interact directly via their LD domains.

To better understand the role of RPN1 in regulating IRE1 activity, we turned to Drosophila melanogaster to explore their interaction *in vivo*. We employed an UAS-Xbp1-HA-GFP reporter line^30^ (provided by Pedro Domingos), which produces GFP upon IRE1-dependent splicing of *Xbp1* mRNA. Combined with GMR-GAL4, this system drives expression specifically in photoreceptor cells, enabling a robust method to track IRE1-mediated splicing activity in vivo. Lines carrying both the reporter and GAL4 driver were crossed with *RPN1* RNAi flies, and IRE1 activity was assessed by measuring GFP fluorescence in the pupal retina using confocal microscopy. Strikingly, RPN1 knockdown resulted in a marked reduction in fluorescence across multiple parameters, suggesting RPN1 is critical for supporting *Xbp1* splicing (Figure 5). Mean fluorescence in visible photoreceptors was significantly lower in *RPN1* knockdowns compared to controls (t(32) = 3.068, p = 0.0044), a trend also observed for integrated density (t(32) = 2.245, p = 0.0318). These results demonstrate that *RPN1* knockdowns significantly impairs IRE1 splicing activity in Drosophila, supporting a functional interaction between RPN1 and IRE1.

**Figure 5.**
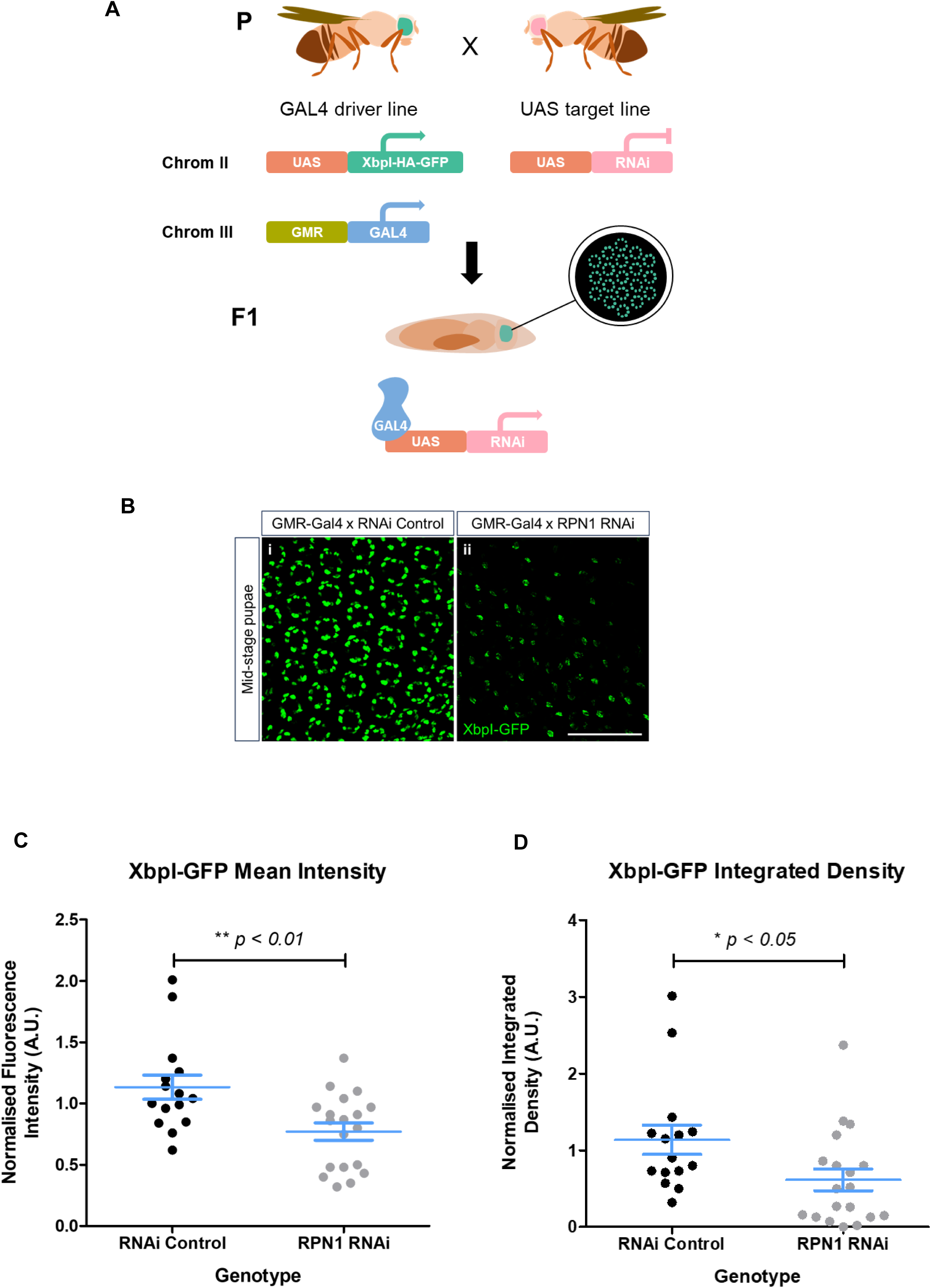
Knocking down RPN1 in vivo causes a reduction in Xbp1-related fluorescence. (**A**) A GAL4-UAS system was used to drive RNAi expression specifically in the Drosophila retina during development. To monitor protein level changes, a *UAS-HA-Xbp1-GFP* transgene was co-expressed, providing a visual readout of spliced Xbp1 post-RNAi induction. (**B**) Representative images of mid-stage (P7-P8) pupal retinas, expressing *UAS-HA-Xbp1-GFP* (green) under *GMR*-GAL4 control. Scale bars represent 50 μm. (**i**) Uniform Xbp1-GFP fluorescence is observed in photoreceptors of control samples. (**ii**) Decreased Xbp1-GFP fluorescence is evident in photoreceptors expressing *RPN1*-RNAi. (**C**) Quantification of mean fluorescence intensity across photoreceptors in all collected samples, shows a significant reduction in RNAi-expressing retinas compared to controls (p < 0.01, Student’s t-test). (**D**) Integrated fluorescence density measurements corroborate the observed (p < 0.05, Student’s t-test). Error bars indicate SEM; statistical significance is denoted (*p < 0.05, **p < 0.01).

## Discussion

In this study we affinity purify and characterise full length IRE1 and PERK protein complexes, and in doing so we overcome a significant technical challenge. Our data suggests a core complex of interacting proteins for both IRE1 and PERK receptors that include RPN1, BiP and Hsp70. An important difference between our current approach and previous studies is that we have purified full-length proteins prior to analysis by mass spectrometry. Furthermore, our samples are derived directly from the separate native gel bands. This enables the examination of distinct IRE1 and PERK complexes, whereas previous studies have used whole cell lysates for analysis without a purification step and have not been able to isolate separate distinct IRE1 and PERK complexes. Furthermore, we have included the investigation of PERK, while previous studies have focused on IRE1 only.

Interestingly, we newly identify RPN1 as an effector of IRE1. RPN1 is observed in all IRE1 and PERK bands within the native gel and directly interacts with IRE1 LD. RPN1 is a subunit of the OST and plays a role in substrate transfer to the catalytic site during N-linked glycosylation^31^. More recently, RPN1 has been shown to be UFMylated in ER-phagy and disrupting this process induces UPR via IRE1 signalling^32^.

In metazoans, XBP1 mRNA is recruited to the ER membrane as part of a nascent chain-ribosome-mRNA complex^23,33^. A defined section of uXBP1 mRNA sequence codes for a hydrophobic stretch that mimics a signal sequence, which is recognised by the SRP and results in the transfer of the nascent chain-ribosome-mRNA complex to the SEC61 translocon^34^. The stability of this complex is supported by a translational pausing mechanism in which the translating peptide forms high affinity interactions in the exit tunnel of the ribosome^23,33^. It is at this stage that IRE1 may engage the translating ribosome and SEC61 to cleave uXBP1 mRNA to generate the spliced version of XBP1 (sXBP1).

Cryo-EM structures have shown that the OST complex is associated with SEC61 whilst it is bound to the ribosome, with the TRAP complex interacting from the opposing side of the SEC61 channel^35^. The SEC61 channel is small and mostly enveloped in the membrane and is dwarfed in size by the ribosome, along with OST and TRAP. The RPN1 subunit does not directly engage with SEC61 as it is on the far side of OST^35^. At this site, RPN1 could provide a stable platform for IRE1 to interact with the mRNA-SEC61-ribosome complex (Figure 6), thereby optimising IRE1 splicing activity. The cryo-EM structures^35^ are consistent with our observation that knockdown of RPN1 inhibits IRE1 from engaging the complex with subsequent loss of splicing activity in the Drosophila retina. Furthermore, our data fits with a model whereby IRE1 is recruited to the ER, possibly interacting with SEC61 and ribosomes so that it can optimally process XBP1 mRNA, whilst the ribosome is translationally paused and receptive to IRE1 splicing^23,33,34^.

**Figure 6.**
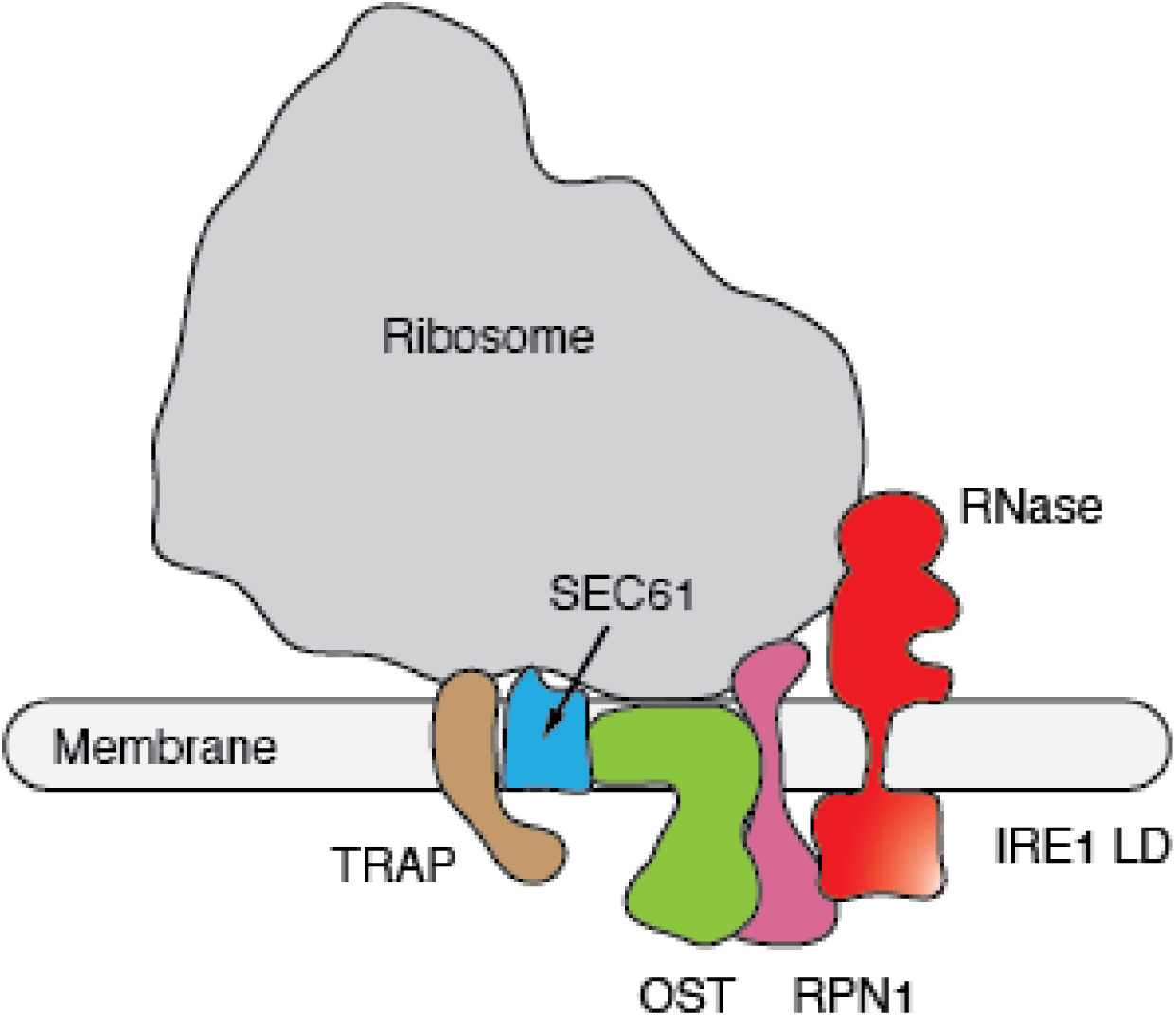
RPN1-SEC61-ribosome complex provides a platform for IRE1 splicing activity. A schematic illustrating the arrangement of SEC61-ribosome-OST complex, with RPN1 domain of OST interacting with IRE1 to provide a platform for IRE1 RNase splicing activity.

The regulation of IRE1 by BiP has been extensively studied. BiP has been shown to act as an inhibitor of IRE1 signalling and its release is thought to be a critical step in IRE1 activation^3,5^. The association of BiP to IRE1 LD would prevent IRE1 interaction with RPN1 and prevent the formation of a productive complex for RNase splicing activity. In high ER stress conditions, BiP release would enable IRE1 to engage RPN1-SEC61-Ribosome complex facilitating the cleavage of XBP1 mRNA. Thus, our data is consistent with the known models of BiP dependent regulation of IRE1^3,5,36^.

Alongside RPN1, both BiP and HSP70 were observed to be present with high scores in all bands that were analysed, suggesting that they form part of a core complex. BiP action has been extensively studied in UPR activation and is known to interact with both IRE1 and PERK to modulate UPR function, although how this occurs is still debated^5,16^. Hsp70 has previously been suggested to form a stable complex with IRE1 and enhance its activity, although the interaction with IRE1 was not influenced by ER stress dependent IRE1 phosphorylation or oligomerisation^37^, similar to our observations with Hsp70.

The expression of IRE1 and PERK in Expi293F suspension cells causes high ER stress equivalent to chemical induction with tunicamycin. In our attempts to mimic a non-ER stressed sample, yet still generate sufficient protein for in vitro analysis, we treated the cell lysate with lambda phosphatase. This successfully reduced phosphorylation on both IRE1 and PERK. However, comparison of the native gel and western blots indicated very little difference between ER stressed and lambda phosphatase treated samples for all the proteins tested except for SEC63, which showed an increase in association in the absence of phosphorylation. The regulation of IRE1 activity by SEC63^22^ has been recently suggested but the influence of phosphorylation was not identified in this previous study^22^. SEC63 is known to be phosphorylated on several positions by CK2 kinase and its association to SEC62 is phosphorylation dependent^38^. In our experiments, the addition of lambda phosphatase to the cell lysate will directly impact other targets including SEC63 and this rather than IRE1 or PERK regulation could explain the phosphorylation linked dissociation from the complex.

The use of lambda phosphatase may not faithfully replicate a non-ER stressed cellular condition; particularly, as the expression of IRE1 and PERK has occurred in stressed cells facilitating BiP dissociation. Subsequent treatment with lambda phosphatase may not be sufficient to return IRE1 and PERK to a non-ER stressed state, thus explaining the lack of difference between lambda phosphatase treated and ER stressed IRE1 and PERK complexes. Experiments conducted in vivo using Drosophila as a model system superseded those conducted in cells. The UAS-Xbp1-HA-GFP reporter under the GAL4 promoter provides a clear phenotype that is specific to IRE1 RNase activity that can be quantitatively measured in vivo, thus providing a powerful means for measuring IRE1 splicing activity.

Interestingly, we identify ITPR3 in only one complex with a high protein score, albeit with relatively low abundance. ITPR3 is an important factor in calcium signalling and its recruitment to mitochondria-ER contact sites is regulated by IRE1. Furthermore, IRE1 influences calcium uptake into the mitochondria via its association with ITPR3^26^. Our data suggests that the interaction of ITPR3 with IRE1 complexes is independent of phosphorylation status of the UPR receptors. Future work will focus on reconstituting individual interacting components to assess their effect on IRE1 and PERK activity and thus will help to clarify their role within such complexes.

The exact oligomeric state of IRE1 and PERK within the various complexes are not readily identifiable as each band has several constituent interacting proteins. The relative abundance of IRE1 and PERK compared to the next most abundant protein varies from being 2:1 to 1:1 possibly suggesting that the UPR receptors are present in monomers, dimers and possibly tetramers. On the other hand, it is plausible that oligomers of IRE1 and PERK interact with oligomers of interacting proteins reflecting the aforementioned abundance ratios, however, this may be unlikely as the size of the complexes derived from the native gel do not allude to very large oligomer species of IRE1 and PERK with interacting proteins in complexes.

In summary, we affinity purify full length IRE1 and PERK and asses the formation of complexes by biochemical and in-vivo analysis. We identify RPN1 as being present in complexes with both IRE1 and PERK. Moreover, knockdown of RPN1 results in a loss of IRE1 RNase activity in the retina of Drosophila. The current study provides a basis for future research to investigate formation and significance of these complexes in UPR signalling and newly identifies RPN1 as an effector of IRE1 RNase activity.

## Methods

### Protein expression in Expi293F cells

Expi293F cells were grown in Expi293^TM^ expression medium (Thermo-Fisher Scientific) using Erlenmeyer tissue culture flasks with vented caps and maintained at 37°C and 8% CO_2_. Cell density and viability were estimated using the Countess^TM^ 3 automated Cell Counter (Thermo-Fisher Scientific). 24h before transfection the cells were diluted to a density of 1.5 x 10^6^ per ml. Transient transfection was performed at a cell density of 2.5 x 10^6^ per ml using polyethylenimine (PEI). 1mg/ml of plasmid DNA was added to PEI at a ratio of 1:2.5 in Opti-MEM® medium (Thermo-Fisher Scientific) and incubated at room temperature for 20 min, then the mixture was added to cells. 24h post transfection, valproic acid was added to the cells at a final concentration of 3.2 mM. The cells were harvested 48h after the transfection by centrifugation at 700g for 5 min with the cell pellet retained. For ER stressed cells, 5µM tunicamycin was added to cells 48h post transfection and cells were harvested 4hrs later.

The cell pellet was resuspended in buffer A that consisted of 10mM Hepes pH7.3, 250mM sucrose, 2mM MgCl_2_ supplemented with protease inhibitor and DNase (sigma). The cells were lysed using a hand homogeniser on ice. The lysed cells were then centrifuged at 2823g at 4°C for 30min to remove the cell debris. The supernatant was collected and centrifuged at high speed of 75000g at 4°C for 1h. The pellet, containing the membrane fraction, was then solubilised in buffer B that consisted of 50mM Hepes pH7.3, 150mM NaCl, 5mM MgAc, 5mM MgCl_2_, 2mM MnCl_2_, 1mM DTT, protease inhibitor (sigma), 1.5% GDN101 (Anatrace) with gentle agitation at 4°C for 2h. The sample was further clarified by centrifugation at 22500rpm for 30mins at 4°C. The supernatant containing the soluble membrane protein was mixed with buffer B without GDN101 (buffer C) in 3 parts to 1 to dilute the GDN101 concentration to below 0.5%.

### FLAG affinity purification

The detergent solubilised sample was applied to 1ml anti-Flag®M2 Affinity resin (Millipore) for 1h at room temperature with gentle agitation, with resin that was pre-equilibrated with buffer C. The sample was then transferred to a gravity flow column at 4°C and the flow through was discarded as the protein was bound to the resin. The sample/resin was washed with 10mls of wash buffer that contained 50mM Hepes pH7.3, 150mM NaCl, 10mM CaCl_2_, 5mM MgCl_2,_ 1mM DTT, protease inhibitor and 0.05% GDN101. Then was washed with 10ml of wash buffer that contained a higher salt concentration of 300mM NaCl to aid removal of unspecific protein bound to resin, before being washed again with 10ml of wash buffer. Protein was eluted by incubating the resin with 4ml of elution buffer which consisted of 50mM Hepes pH7.3, 150mM NaCl, 10mM CaCl_2_, 5mM MgCl_2_, 1mM DTT, protease inhibitor, 0.05% GDN101, 250µg/ml Flag peptide (sigma), for 15mins at 4°C. The elution step was repeated for another 3 times to ensure all protein was eluted. The protein was concentrated to a volume of 100ul using a single amicon 100K MW cut off at 4°C.

For lambda phosphatase treated sample, 500µl of lambda phosphatase at a concentration of 40mg/ml was added to the detergent solubilised sample, which had been split into two parts, during the 1hr incubation with FLAG resin. The other part was treated separately and acted as control measurement.

### Western blot analysis

Samples were mixed in a 1:1 ratio with Laemmli buffer (Sigma) and boiled at 95°C. Proteins were separated using 4-15% SDS PAGE gel (Biorad). Proteins were transferred from gel to nitrocellulose membrane using iBlot2. The membrane was blocked in TPBS buffer with 5% dried milk for 1hr at room temperature and then incubated with primary antibody in TPBS + 2% milk for 1hr at room temperature. The membrane was washed 3 times with TPBS for 5mins and incubated with secondary antibody in TPBS + 2% milk for 1hr then washed again. Blots were visualised using Pierce ECL western blot (thermo-scientific) on a ChemiDoc imaging system (biorad).

The primary antibodies were purchased from abcam for BiP (ab108613), SEC63 (ab685550), GCN1 (ab86139), SEC61A (ab183046), ITPR3 (ab264283), RPS6 (ab70227), RPL18 (ab 241988), IRE1 (ab227245), IRE1 phosphorylated (ab48187), PERK (ab65142); from Invitrogen for Hsp70 (PA5-11473) and PERK phosphorylated (PA5-40294); from Sigma for RPN1 (WH0006184-100UG) and FLAG (SAB4301135-100UG). Anti-rabbit and anti-mouse HRP linked secondary antibodies were purchased from cell signalling technology.

### Native gel analysis

Native gel reagents were purchased from Thermo scientific. FLAG purified protein was added to NativePAGE sample buffer and G-250 sample additive and made up to 15µl volume with deionised water. Samples were added to native Bis-Tris 3-12% gel with NativeMark unstained proteins standard. Dark blue cathode with 0.02% (w/v) Coomassie G-250 and anode buffer were carefully applied to fill the chambers of mini cell. Gel was run for 4C for 30mins at 150V, after which cathode buffer was replaced with light blue buffer with 0.002% Coomassie G-250 and continued at 250V for 150mins. Gels were visualised using Coomassie G-250 following the manufactures protocol.

### Mass spectrometry

The excised gel bands were diced into 1 mm cubes, reduced, alkylated and digested with trypsin. 20 ul of peptides were extracted and a portion between 1-5 ul – dependant on the strength of the Coomassie staining – was analysed by LCMSMS. Analysis was carried out on a Thermo scientific Ultimate3000 at 300 nl/min with a reverse phase linear gradient of water with 0.1 % formic acid to acetonitrile with 0.1 % formic acid, over 1 hour. The column used was a Thermo scientific pepmap easyspey column 75 um x 15 cm and the eluting peptides sprayed directly into the Thermo scientific Fusion Lumos mass spectrometer. The mass spectrometer was operated in data dependant mode scanning MS from 375-1500 m/z on the orbitrap at 60000 resolution. The top 10 most intense peptides with charge states between 2-5+ were selected for HCD fragmentation and MSMS data collected from 140-2000 m/z on the iontrap. The raw data was analysed in Mascot 2.6 (MatrixScience) using the msconvert data extraction script. Data was searched against the SwissProt database of proteins restricted to human only proteins (563,082 protein sequences downloaded Apr 2020) using trypsin as digestion enzyme, carbamidomethyl as fixed modification and oxidation of methionine as variable modification with 20 ppm and 0.6 Da search tolerances on the MS and MSMS searches respectively. The protein score and the relative abundance (emPAI value) was used to assess selected hits. The protein score is derived from the ion score, which is calculated based on the probability, P, that the match between experimental data and database sequence is a random event. The score is reported as −10Log(p). A threshold of p<0.05 indicate that protein scores of above >35 relates to identity or extensive homology.

### Fly Husbandry

All *Drosophila* crosses were raised at 25°C in a 12hr light/dark cycle, on standard cornmeal media prepared in-house. Experimental flies were generated by crossing *w-; UAS-XbpI-HA-GFP/CyO, DGY; GMR-Gal4/TM6b, DGY* virgin females to *KK RNAi control* males (VDRC stock #60101) or *RPN1 KK RNAi* males (VDRC stock #107778). F1 offspring were collected and aged appropriately before dissection.

### Immunohistochemistry

For dissections of pupal retinas, late third-instar larvae were selected based on the inheritance of genetic markers and maintained at 25°C for 50 hours. Pupal brains were dissected in 1X PBS following the method outlined by Coelho et al. (2013)^39^. After dissection, samples were fixed in 4% formaldehyde in 1X PBS for 40 minutes at room temperature, followed by three washes with 0.3% Triton X-100 in 1X PBS (PBST). Retinas were isolated and mounted in Vectashield antifade medium to minimize photobleaching and subsequently analysed using a Zeiss LSM 510 confocal microscope.

### Image Analysis

Fluorescence quantification and particle property analysis were conducted using ImageJ software (version 1.54f, National Institutes of Health, USA). Confocal images of pupal retinas, expressing endogenous GFP, were processed by Z-projection and analysed for mean gray value and integrated density (IntDen) using a custom macro (Supplementary Data 5). Statistical analyses were carried out in GraphPad Prism (version 5.0). To account for inter-experimental variability, datasets were normalised to relevant means, with outliers objectively identified and excluded where appropriate. Group differences were evaluated using an unpaired t-test, and significance annotated in graph figures.

## Data availability

Data supporting this study are included within the article and supplementary/source material.

## Acknowledgements

We are grateful to Prof. Pedro Domingos and Fatima Cairrao (ITQB-UNL, Portugal) for providing the UAS-XBP1-UA-GFP construct, fly line and for their helpful advice. We would like to P. Nowak and Carina Rassam for assistance with protein expression and purification. Also, we would like to thank the mass spectrometry and proteomics facility, University of St Andrews, UK. This work was funded by CRUK fellowship to (grant no. C33269/A20752 and C33269/A23215) and MRC grant (MR/Y012224/1) awarded to MMUA.

## Author contribution

NL, FS, YBK - conceptualised and designed experiments, conducted data acquisition and research investigation, analysed and interpretated the data, and reviewed draft manuscript. CSC - conceptualised and designed experiments, conducted data acquisition and research investigation, analysed and interpretated the data. CA, - data acquisition, analysis and interpretation. TS - conceptualised and designed experiment, analysed and interpreted data, and drafted manuscript. MMUA – conceptualised and designed experiment, analysed and interpreted data, drafted manuscript, supervised project and obtained funding.

## Competing interests

The authors have no competing interests

---

**Supplementary Data 1.** Mass spectrometry-based Protein ID of bands excised from SDS PAGE gel. The protein scores are based on ion scores with a value of >35 indicating identity or extensive homology.

|  | Score | Mass<br>FL protein | peptide matches | unique sequence |
| --- | --- | --- | --- | --- |
| PERK | 326 | 126.1 | 15 | 14 |
| IRE1 | 227 | 110.5 | 19 | 13 |
| SEC63 | 115 | 88.3 | 10 | 7 |
| Hsp90 | 240 | 83.5 | 15 | 14 |
| BiP | 179 | 72.4 | 8 | 8 |
| Hsp70 | 127 | 70.2 | 5 | 4 |
| RPL3/4 | 162 | 47.9 | 11 | 9 |
| SEC61 | 203 | 52.6 | 16 | 10 |

**Supplemental Data 2.**
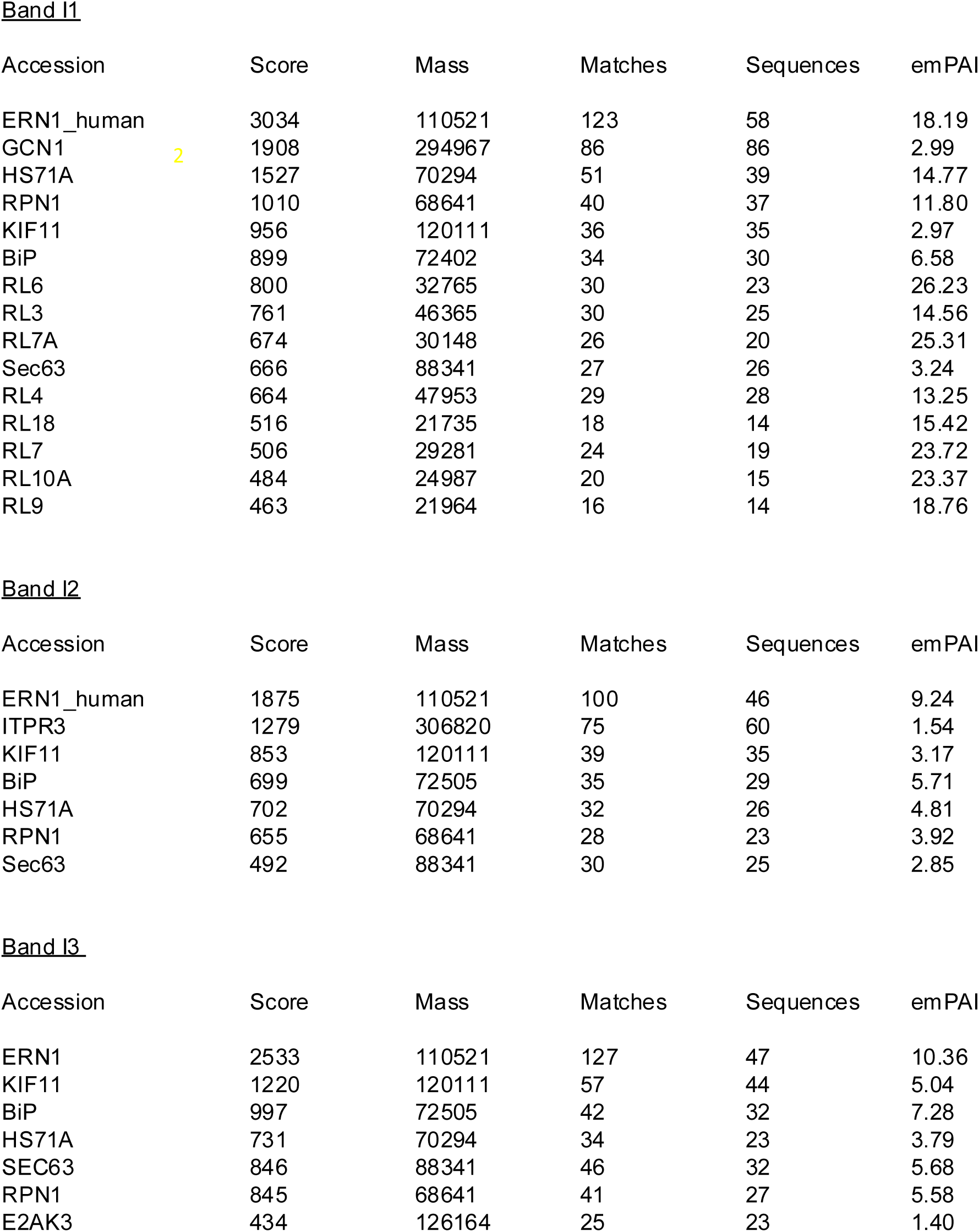
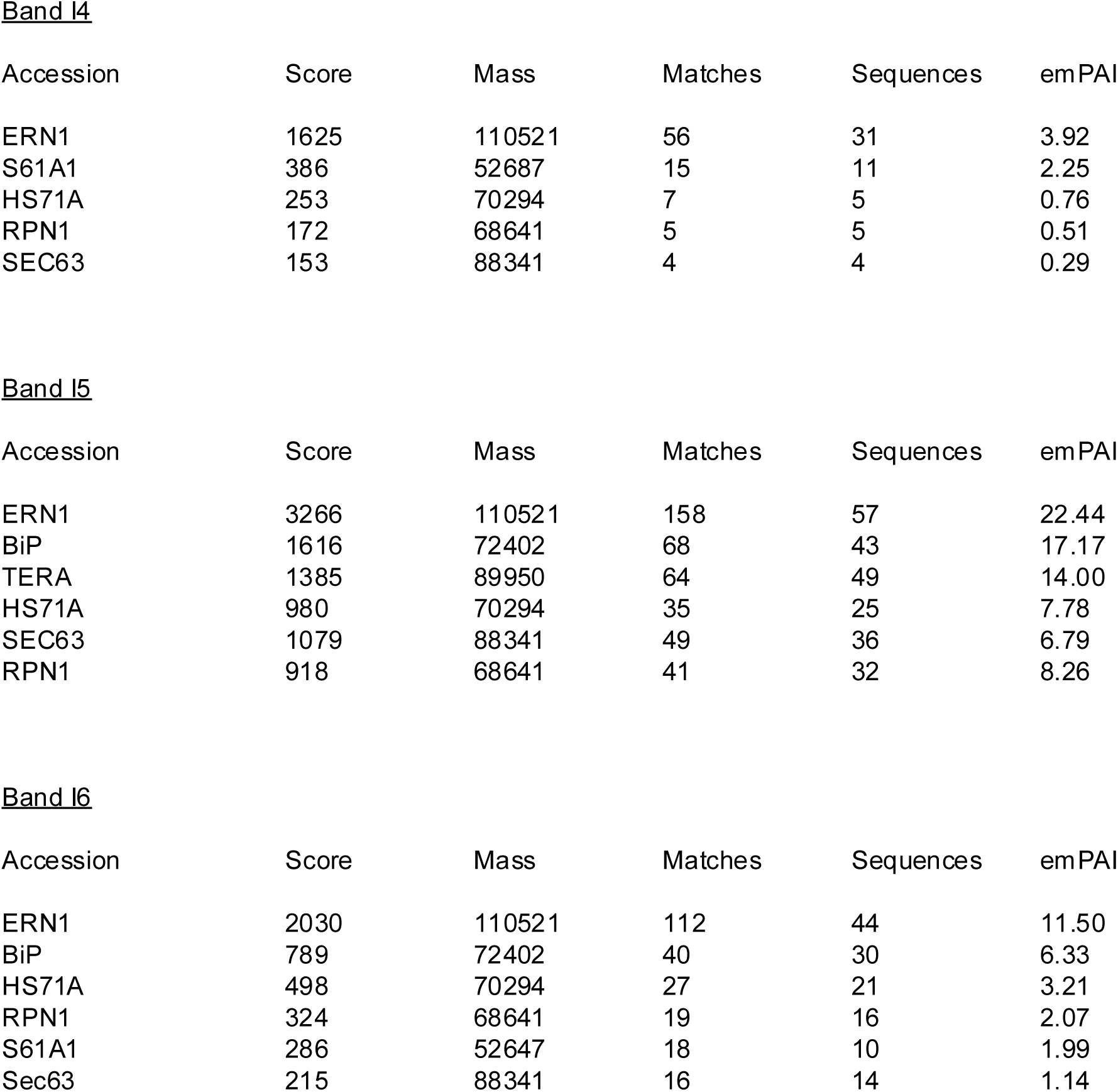
Mascot analysis of ESI MS/MS data for each of the excised bands isolated from the native gel performed on IRE1 full length protein. The protein that are listed are selected based on high protein score cut off point relative to other proteins from the Mascot output. The protein scores are based on ion scores with score value greater >35 indicating identity or extensive homology. The emPAI measures the relative abundance of proteins within each sample.

**Supplemental Data 3.** Mascot analysis of LCMS/MS data for each of the excised bands isolated from the native gel performed on PERK full length protein. The protein that are listed are selected based on high protein score cut off point relative to other proteins from the mascot output. The protein scores are based on ion scores with score value greater >35 indicating identity or extensive homology. The emPAI measures the relative abundance of proteins within each sample.

Band P1 PERK
| Accession | Score | Mass | Matches | Sequences | emPAI |
| --- | --- | --- | --- | --- | --- |
| E2AK3 | 2626 | 126164 | 123 | 65 | 11.83 |
| HS71A | 937 | 70294 | 44 | 31 | 7.89 |
| BiP | 460 | 72402 | 19 | 15 | 1.70 |
| RL4 | 691 | 47953 | 28 | 21 | 8.93 |
| RL3 | 674 | 46365 | 27 | 20 | 6.86 |
| RL7A | 621 | 30148 | 25 | 19 | 19.06 |
| RL18 | 601 | 21636 | 20 | 10 | 6.19 |
| RPN1 | 508 | 68641 | 24 | 19 | 2.78 |
| RL6 | 468 | 32765 | 23 | 16 | 9.28 |
| DNJA3 | 403 | 53083 | 19 | 14 | 2.54 |
| GCN1 | 320 | 294967 | 22 | 19 | 0.36 |
| KIF11 | 316 | 120111 | 17 | 16 | 0.90 |

Band P2
| Accession | Score | Mass | Matches | Sequences | emPAI |
| --- | --- | --- | --- | --- | --- |
| E2AK3 | 2974 | 126164 | 140 | 68 | 13.54 |
| HS71A | 872 | 70294 | 46 | 32 | 8.30 |
| BiP | 646 | 72402 | 28 | 22 | 3.23 |
| RPN1 | 822 | 68641 | 44 | 36 | 11.08 |
| GCN1 | 489 | 294967 | 35 | 33 | 0.70 |
| KIF11 | 455 | 120111 | 22 | 20 | 1.21 |

Band P3
| Accession | Score | Mass | Matches | Sequences | emPAI |
| --- | --- | --- | --- | --- | --- |
| E2AK3 | 2727 | 126164 | 136 | 63 | 10.47 |
| RPN1 | 988 | 68641 | 46 | 33 | 9.41 |
| HS71A | 938 | 70294 | 47 | 30 | 7.05 |
| ECHA | 804 | 83688 | 39 | 30 | 4.77 |
| BiP | 760 | 72402 | 35 | 26 | 4.45 |

Band P4 (9)
| Accession | Score | Mass | Matches | Sequences | emPAI |
| --- | --- | --- | --- | --- | --- |
| E2AK3 | 3787 | 126164 | 170 | 69 | 14.49 |
| PRKDC | 1227 | 473749 | 72 | 67 | 0.96 |
| ECHA | 1121 | 83633 | 49 | 35 | 8.09 |
| HS71A | 1099 | 70294 | 45 | 33 | 8.86 |
| BiP | 1044 | 72402 | 45 | 36 | 9.51 |
| KIF11 | 870 | 120111 | 36 | 31 | 2.53 |
| RPN1 | 745 | 68641 | 35 | 30 | 6.91 |

Band P5 (10)
| Accession | Score | Mass | Matches | Sequences | emPAI |
| --- | --- | --- | --- | --- | --- |
| E2AK3 | 3350 | 126164 | 158 | 71 | 14.47 |
| GCN1 | 1275 | 294967 | 75 | 29 | 2.09 |
| HS71A | 1132 | 70294 | 48 | 33 | 8.85 |
| BiP | 1009 | 72402 | 41 | 33 | 8.22 |
| RPN1 | 971 | 68641 | 37 | 30 | 6.90 |

Band P6 (11)
| Accession | Score | Mass | Matches | Sequences | emPAI |
| --- | --- | --- | --- | --- | --- |
| E2AK3 | 3437 | 126164 | 130 | 61 | 10.91 |
| TERA | 903 | 89950 | 45 | 36 | 6.79 |
| HS71A | 803 | 70294 | 43 | 30 | 7.06 |
| BiP | 612 | 72402 | 29 | 25 | 4.12 |
| RPN1 | 517 | 68641 | 29 | 22 | 3.56 |

**Supplemental Data 4.**
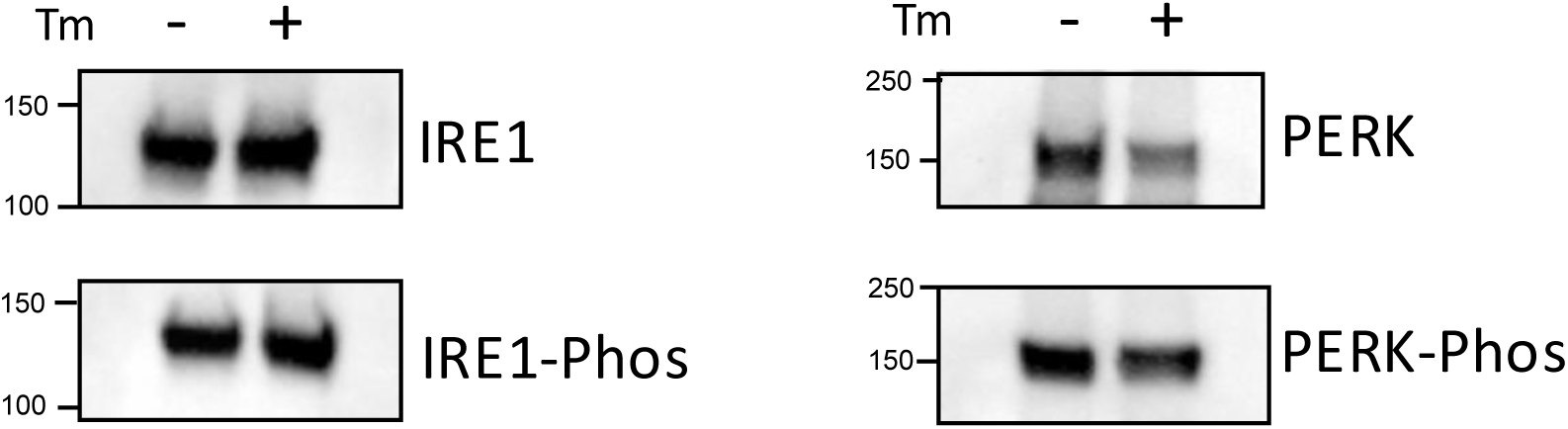
Expi293F cells expressing IRE1 and PERK were subjected to 5μm tunicamycin for 4hrs and then harvested. The proteins were FLAG affinity purified from cells and their phosphorylation status were assessed by western blot using phospho specific antibodies for IRE1 and PERK.

**Supplemental Data 5.**
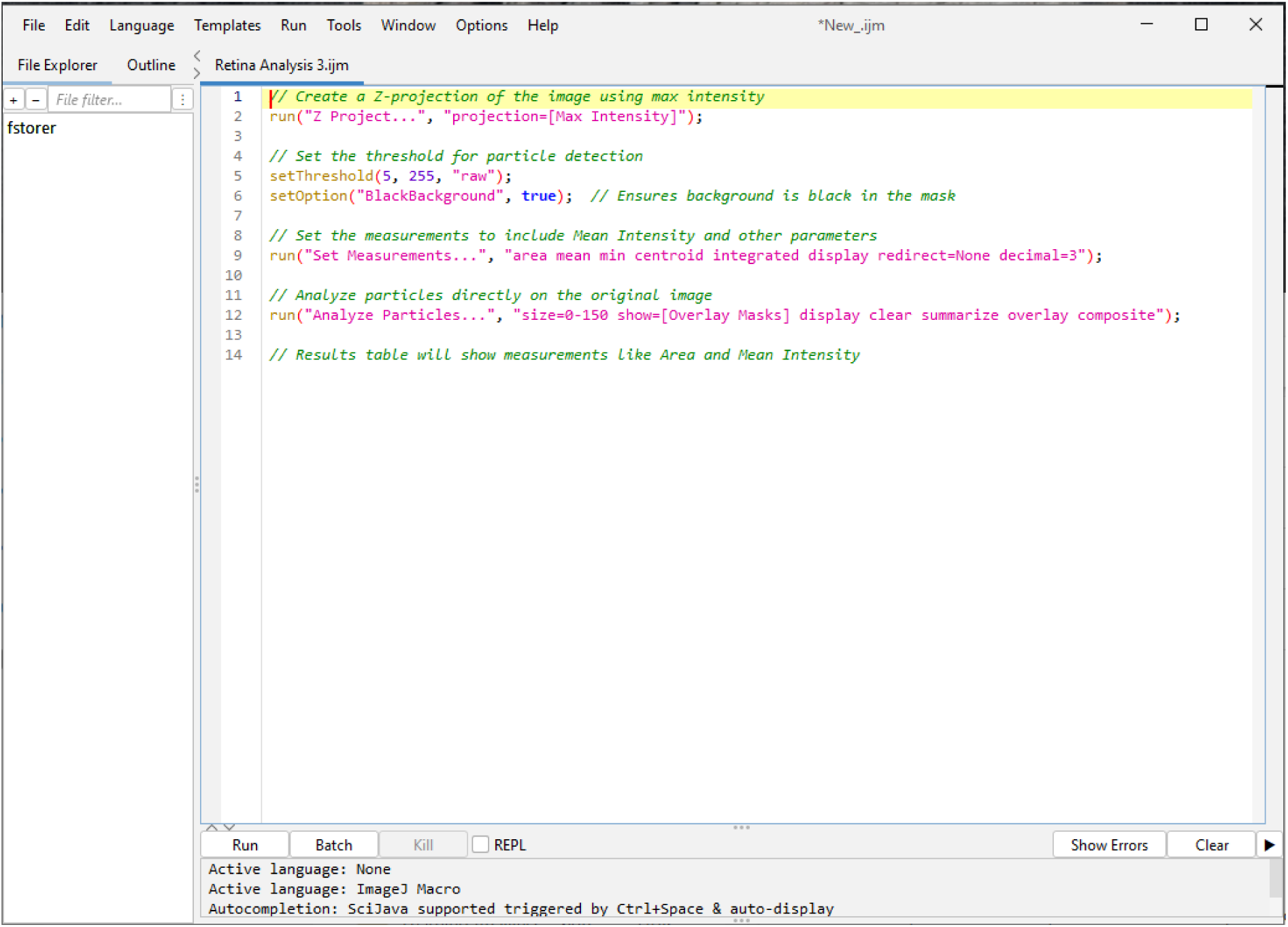
Confocal images of pupal retinas, expressing endogenous GFP, were processed by Z-projection and analysed for mean gray value and integrated density (IntDen) using the custom macro listed.

## Notes

### Competing Interest Statement

The authors have declared no competing interest.

